# Deriving longitudinal tumor phylogenies from single-cell sequencing data

**DOI:** 10.1101/2025.08.06.668977

**Authors:** Akhil Jakatdar, Palash Sashittal, Benjamin J. Raphael

## Abstract

Tumors evolve over time and in response to treatment, leading to changes in the proportions of clones within the tumor. Single-cell sequencing of tumor samples from multiple timepoints enables the reconstruction of clonal evolution and tracking of temporal changes in tumor composition. However, the high rates of missing data in single-cell sequencing complicate evolutionary analyses, and can lead to implausible conclusions, such as the reappearance of extinct clones. We introduce Phyllochron, an algorithm that builds a *longitudinal phylogeny* from cancer cells sequenced from multiple timepoints and evaluates whether the constraints on clonal proportions imposed by longitudinal sampling are well supported by the data. Phyllochron relies on a novel mathematical formulation of a longitudinal perfect phylogeny, an extension of the perfect phylogeny model that is widely used in cancer evolution. We show that Phyllochron outperforms existing phylogeny inference methods on simulated single-cell sequencing data. Applied to longitudinal single-cell DNA sequencing data from an acute myeloid leukemia (AML) patient, Phyllochron constructs a longitudinal phylogeny containing rare cancer clones that persist through multiple cycles of targeted drug treatment, a crucial finding missed by existing phylogeny inference methods. The Phyllochron statistical test supports the presence of these rare clones.

## 1 Introduction

Cancer is driven largely by the accumulation of somatic mutations in a population of cells. This process results in a heterogeneous tumor made up of subpopulations of cells, called *clones*, with distinct sets of somatic mutations [1], which evolve over time and in response to treatment. For example, clones that were initially present in small proportions before treatment may become dominant and lead to relapse or metastasis. Tracking the evolution and clonal composition of a tumor through time is crucial for understanding relapse and resistance [2–8].

Single-cell DNA sequencing (scDNA-seq) measures somatic mutations from hundreds or thousands of cells from samples of the same tumor and enables the tracking of the evolutionary history of the tumor. However, reconstruction of the evolution and clonal composition of a tumor from such scDNA-seq data is challenging due to errors and missing data. For example, targeted scDNA-seq technologies such as MissionBio Tapestri measure single nucleotide variants (SNVs) with high fidelity but suffer from high rates of missing data (up to 20% [9]) due to technical limitations such as amplification bias and allelic dropout [10–13]. Specialized methods for inferring tumor phylogenies from scDNA-seq employ evolutionary models to correct errors and impute missing mutations in the data [14–27]. However, due to uncertainty in imputation and error correction, there are often multiple plausible tumor phylogenies equally supported by the data [19, 22, 25].

Measuring an evolutionary process at multiple timepoints, known as *longitudinal sampling*, provides additional information for phylogenetic tree inference since longitudinal samples have a shared evolutionary history. Specifically, longitudinal sampling measures ancestral cells, or internal nodes in the phylogeny, while single timepoint sampling measures only extant cells, or leaves in the phylogeny. In *stratocladistics* information about ancestral species – such as the morphology of fossils or the age of the strata in which they were found – are used when inferring evolutionary relationships between species [28–30]. Subsequently, several methods have demonstrated the benefits of using the temporal signal in longitudinal data for inferring viral phylogenies from serially sampled viral sequences [31–34] and tumor phylogenies from longitudinal bulk DNA sequencing data [35].

Existing methods for reconstructing tumor phylogenies from longitudinal scDNA-seq data are limited in their ability to exploit the temporal information. For example, [36] treats each timepoint as an independent sample and applies phylogeny inference methods originally developed for single timepoint data, e.g. SCITE [14] and COMPASS [27], to these samples. Interestingly, this approach sometimes produces phylogenies where cancer clones disappear at one timepoint and subsequently reappear at a later timepoint, which is an unlikely scenario if the tumor heterogeneity is well-sampled using current high-throughput scDNA-seq technologies [13, 37]. More recently, methods such as LACE [38] and scLongTree [39] were developed specifically for longitudinal scDNA-seq data. However, LACE does not explicitly model the temporal constraints imposed by longitudinal sampling on the phylogenies, and neither method evaluates whether such constraints are supported by the longitudinal scDNA-seq data of a tumor.

We introduce Phyllochron, an algorithm to reconstruct a phylogeny from longitudinal scDNA-seq data and evaluate whether the constraints imposed by reasonable assumptions on longitudinal sampling are well supported by the data. Phyllochron integrates an algorithm to infer longitudinal perfect phylogenies with a statistical test to evaluate whether the observed read counts of mutations are better fit by longitudinal sampling. The framework is built on the *longitudinal perfect phylogeny* model, an extension of the perfect phylogeny model [40], which has been widely applied in cancer phylogenetics [14, 15, 18, 19, 35]. We formalize the problem of inferring longitudinal perfect phylogenies under a range of sequencing error models and analyze its computational complexity. We show that Phyllochron reconstructs more accurate tumor phylogenies compared to existing methods on simulated data. On longitudinal targeted scDNA-seq data of an acute myeloid leukemia (AML) patient, we find that temporally inconsistent events inferred by other methods, such as the same clone disappearing and reappearing during treatment or a clone appearing before its ancestor, are not well supported by the data. In contrast, Phyllochron finds evidence that all these clones are present before and after treatment of the tumor, a crucial finding that is missed by existing methods.

## 2 Results

### 2.1 Longitudinal Sequencing and Longitudinal Phylogenies

Longitudinal sequencing experiments yield sequential measurements of a tumor across multiple timepoints. Suppose we perform longitudinal sequencing of a tumor at *k* timepoints, measuring *m* somatic mutations from *n* total cells. We represent these measurements with an *n* × *m* binary mutation matrix *A*, where entry *a*_*i,j*_ = 1 if mutation *j* is present in cell *i* and *a*_*i,j*_ = 0 if the mutation is absent. The sampling time of the cells is encoded by a timepoint vector ***τ***, where each entry *τ*_*i*_ ∈ {1,…, *k*} is the timepoint at which cell *i* was sampled.

We represent the evolutionary history of cells measured across multiple timepoints by a longitudinal phylogeny *T*. A longitudinal phylogeny *T* = (*V* (*T*), *E*(*T*)) is a rooted tree in which vertices *V* (*T*) = {1,…, *n*} represent cells and edges *E*(*T*) connect parental cells to their descendants. Edges are labeled by mutation(s) that distinguish descendants from parental cells. In contrast to standard phylogenies, where only the leaves correspond to observed cells, longitudinal phylogenies may include labeled internal nodes corresponding to cells that were measured at earlier timepoints. That is, some ancestral cells in the tumor may be directly observed and thus labeled in the phylogeny. We label each observed cell *i* in the longitudinal phylogeny with its mutation vector **a**_*i*_ ∈ {0, 1}^*m*^ and its sampling time *τ*_*i*_. Hence, a longitudinal phylogeny is a partially vertex-labeled tree where all the leaves and some internal vertices are labeled by a mutation vector and a timepoint. A key property of longitudinal phylogenies is *temporal consistency*, which requires that an ancestor must be sampled no later than its descendants. Formally, if a labeled vertex *u* lies on the path from the root to another labeled vertex *v*, then *τ*_*u*_ < *τ*_*v*_. Temporal consistency ensures that the sequence of sampling times along any path from root to leaf is strictly increasing. This constraint arises directly from the structure of the data and does not rely on any assumptions about the sampling process. Note that longitudinal phylogenies are distinct from *timed phylogenies*, which have previously been used to model temporal sequencing data [41–43], as discussed in Methods section 4.1.

Since current single-cell sequencing technologies are destructive and remove cells from the population for sequencing, we introduce an additional assumption about the relationship between cells sampled at different timepoints. Specifically, the destructive nature of sequencing means that the descendants of a sequenced cell are not measured at later timepoints. We assume that for every sequenced cell at time *i*, there exists a cell with an identical mutation profile (up to the measured mutations; e.g., from the same cancer clone) that remains in the population and continues to divide. We formalize this as the *ancestral sampling assumption*: for each cell sampled at time *t* > 1, an ancestor was sampled at time *t* − 1. Thus, under the ancestral sampling assumption, the timepoints of the labeled vertices along every path from the root to a leaf in a longitudinal phylogeny must appear in a consecutive, strictly increasing order from 1 to *k*′ for some timepoint *k*′. While idealized, this assumption becomes increasingly plausible with higher-resolution sampling and tumors composed of stable clones. Moreover, in Section 2.4, we show that this assumption is often less restrictive than those made by existing methods, and we evaluate its plausibility using a hypothesis test in Section 4.5.

Longitudinal phylogenies are compatible with a wide range of evolutionary models. The structure of a longitudinal phylogeny is determined by both the evolutionary process and the temporal constraints imposed by sampling. As such, any model that defines how mutations accumulate over time can be incorporated, including, but not limited to, the *perfect phylogeny model* [44], the *Dollo model* [45], the *constrained Dollo model* [16] and the *finite sites model* [40]. Our formulation of longitudinal phylogenies provides a foundation for integrating diverse models of tumor evolution with time-resolved single-cell data.

### 2.2 Phyllochron: an algorithm to infer and evaluate support of longitudinal phylogenies

We introduce Phyllochron, an algorithm to infer and evaluate support for longitudinal phylogenies from longitudinal single-cell DNA sequencing (scDNA-seq) data. Our algorithm is composed of two parts: (1) an algorithm to infer longitudinal phylogenies under the perfect phylogeny model; and (2) a statistical test that evaluates whether the inferred longitudinal phylogeny is supported by the data (Figure 1).

**Fig. 1.**
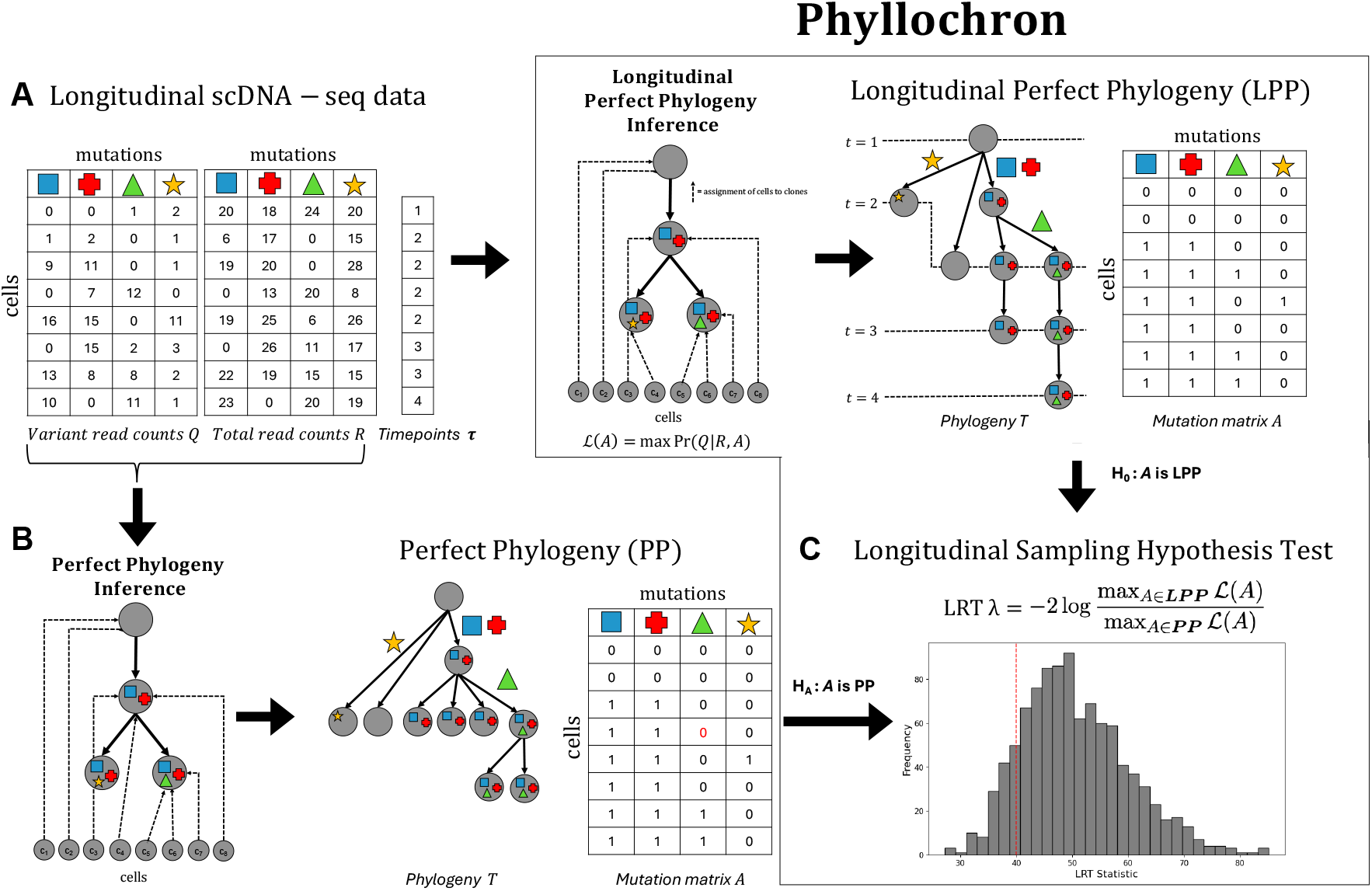
Overview of Phyllochron. (A) Phyllochron takes read count matrices *Q* and *R* and timepoints *τ* from a longitudinal single-cell sequencing experiment as input, and constructs a maximum likelihood longitudinal perfect phylogeny *T* and corresponding mutation matrix *A* under a probabilistic read count error model. (B) The workflow of current maximum likelihood perfect phylogeny methods that do not explicitly model longitudinal data. (C) The Phyllochron hypothesis test computes the likelihood ratio test (LRT) statistic *λ* from a maximum likelihood longitudinal perfect phylogeny (LPP) and a perfect phylogeny (PP) solution. Phyllochron compares the *λ* value for the data (red line) to an empirical distribution of LRT values obtained on permuted datasets.

Phyllochron derives a longitudinal phylogeny under the perfect phylogeny model using a probabilistic model for the sequencing data. The perfect phylogeny model [44] is a widely used model for cancer evolution [14, 19, 46, 35] that assumes that each mutation occurs only once and is not subsequently lost. We define a *longitudinal perfect phylogeny* as a longitudinal phylogeny whose mutation matrix satisfies the perfect phylogeny assumptions. Specifically, a longitudinal perfect phylogeny is a rooted tree in which each mutation labels exactly one edge, and every path from the root to a leaf corresponds to a sequence of cells sampled at timepoints that appear in consecutive and increasing order (see Section 4.1 for a formal definition).

In scDNA-seq, we do not observe the binary mutation matrix *A* directly but instead measure the number of reference and variant reads at the mutation loci in each sequenced cell. Specifically, we observe an *n* × *m* variant read count matrix *Q* = [*q*_*i,j*_], where *q*_*i,j*_ is the number of reads supporting the variant allele, and an *n* × *m* total read count matrix *R* = [*r*_*i,j*_], where *r*_*i,j*_ is the total number of reads covering mutation *j* in cell *i*. We model the observed read counts using a beta-binomial distribution, similar to previous works [25, 22, 17, 27] (details in Section C.1). Given matrices *Q* and *R* and a timepoint vector ***τ***, our goal is to infer a mutation matrix *A* that maximizes the likelihood Pr(*Q* | *R, A*) of observing the read counts and admits a longitudinal perfect phylogeny *T* with respect to the timepoints ***τ***. We formalize this as follows:

#### Longitudinal Perfect Phylogeny from Read Counts (LPP-RC)

*Given a variant read count matrix Q, total read count matrix R, and a timepoint vector* ***τ***, *find a mutation matrix A and longitudinal perfect phylogeny T such that (i) A maximizes the likelihood* Pr(*Q* | *R, A*) *and (ii) A and* ***τ*** *admit T*.

We show that the LPP-RC problem is NP-hard (Section 4.2) and implement an algorithm in Phyllochron to solve this problem. The algorithm relies on a combinatorial characterization of longitudinal perfect phylogenies (Section 4.3). Using this characterization, we derive a mixed integer linear program (MILP) to solve a variant of the LPP-RC problem where the *clone tree* describing the evolutionary relationships between the cancer clones is given. Specifically, the MILP assigns cells to clones such that the resulting mutation matrix *A* maximizes the likelihood Pr(*Q R, A*) such that *A* admits a longitudinal perfect phylogeny. Phyllochron solves the LPP-RC problem using an iterative hill-climbing approach to search the space of clone trees, where the MILP is used to evaluate the candidate clone trees at each iteration (details in Section 4.4.2).

The longitudinal constraints derived from the ancestral sampling assumption are enforced under the longitudinal perfect phylogeny model, and so we need a rigorous test to evaluate the validity of this assumption. Here, we introduce a hypothesis test to assess the validity of longitudinal phylogenies on a given dataset. Specifically, we compare two nested models:

- Null hypothesis: The observed variant read counts *Q* are generated from a longitudinal perfect phylogeny tree.
- Alternative hypothesis: The observed variant read counts *Q* are generated from a perfect phylogeny tree.

These models are nested because every longitudinal perfect phylogeny is also a perfect phylogeny. We employ a likelihood ratio test to evaluate these hypotheses by computing maximum likelihood trees under both the null and alternative hypotheses for a fixed clone tree. Specifically, we compute the likelihood ratio test (LRT) statistic 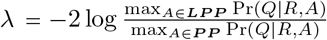. We assess the statistical significance of the log-likelihood ratio using a permutation test, as described in Section 4.5.

### 2.3 Simulated data

We compare Phyllochron with ConDoR*^1^ [25], COMPASS [27], scLongTree [39], and LACE [38] on simulated data. We simulate longitudinal perfect phylogenies with 10 mutations on simulated instances with 3, 4 and 5 timepoints with varying rates of missing data (0.01, 0.10, and 0.20). We encountered issues with scalability and numerical stability with the longitudinal methods scLongTree and LACE (Supplementary Section C.5), and thus compared these methods to Phyllochron only on a smaller simulation with *n* = 100 cells. We compared Phyllochron with ConDoR* and COMPASS on a larger simulation with *n* = 1000 cells. Details of all method parameters and a runtime analysis of all methods are provided in Supplementary Section C.5 and Supplementary Figure S4, respectively.

We evaluate the accuracy of each method by computing the *clone lifespan error*, the number of longitudinal violations and the *mutation matrix error* [16, 22, 25]. The clone lifespan error evaluates the accuracy in reconstruction of the lifespan, or set of timepoints at which cells from the clone are present, for all clones in the tumor. Formally, the clone lifespan error is the average Jaccard distance over the *m* clones between the ground truth and inferred set of timepoints where the clone is present. The number of longitudinal violations is the number of clones in the inferred phylogeny that have either a temporal inconsistency or an ancestral sampling violation (details in Section 4.6). Lastly, the mutation matrix error is the normalized *L*_1_ norm between the ground-truth mutation matrix *A* and the inferred mutation matrix *Â*. Phyllochron, ConDoR*, and COMPASS take the read count matrices as input, while scLongTree and LACE take an observed mutation matrix *A*′ as input. We derive *A*′ from the simulated read counts as described in Supplementary Section C.

On the small simulations consisting of *n* = 100 cells, Phyllochron outperforms competing longitudinal methods on all three metrics across all simulation parameters (Figures 2 and S3). For instance, on the simulations with *k* = 5 timepoints and a missing data rate of *d* = 0.2, Phyllochron achieves the lowest median clone lifespan error (*δ* = 0.030) and mutation matrix error (*ϵ* = 0.012), outperforming scLongTree (*δ* = 0.12, *ϵ* = 0.027) and LACE (*δ* = 0.136, *ϵ* = 0.206) (Figure 2(A,C)). Both Phyllochron and scLongTree phylogenies have no longitudinal violations, while LACE has longitudinal violations in 26 of the 30 simulated phylogenies (Figure 2(B)). Although scLongTree infers phylogenies without longitudinal violations, it has higher mutation matrix error and clone lifespan error than Phyllochron, likely because scLongTree uses various heuristics while Phyllochron is based on a rigorous definition of a longitudinal phylogeny.

**Fig. 2.**
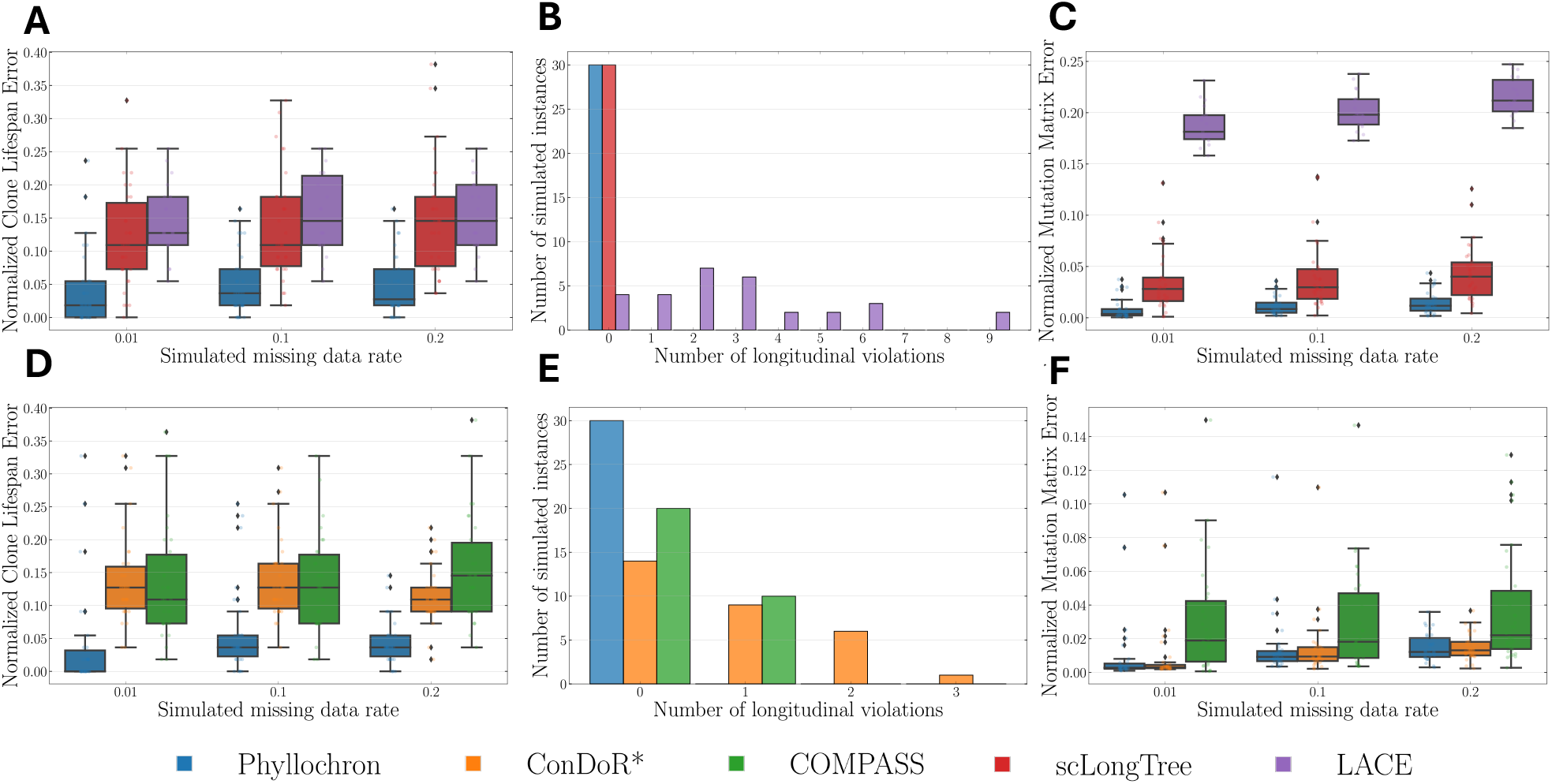
Phyllochron outperforms existing methods in accurately inferring the phylogeny and clone lifespans on simulated longitudinal sequencing data. (A) The normalized clone lifespan error of Phyllochron and longitudinal methods scLongTree and LACE. (B) The number of longitudinal violations in phylogenies inferred by Phyllochron, scLongTree and LACE. (C) The normalized mutation matrix error of the mutation matrices imputed by Phyllochron, scLongTree and LACE. (D) The normalized clone lifespan error of Phyllochron and non-longitudinal methods ConDoR* and COMPASS. (E) The number of longitudinal violations in phylogenies inferred by Phyllochron, ConDoR* and COMPASS. (F) The normalized mutation matrix error of Phyllochron, ConDoR* and COMPASS. All methods are evaluated against the simulated ground truth with *k* = 5 timepoints under varying missing data rates (see Supplementary Section C). Box plots show the median and the interquartile range (IQR), and the whiskers denote the lowest and highest values within 1.5 times the IQR from the first and third quartiles, respectively. Histograms show the distribution of the number of longitudinal violations for each method across all 30 replicates.

On larger simulations with *n* = 1000 cells, Phyllochron significantly outperforms both ConDoR* and COMPASS on clone lifespan error and the number of longitudinal violations (Figure 2(D,E)). On mutation matrix error, Phyllochron performs competitively with ConDoR* and outperforms COMPASS (Figure 2(F)). On simulations with *k* = 5 timepoints and missing data rate *d* = 0.2, Phyllochron achieves the best clone lifespan error (*δ* = 0.061) while maintaining a low mutation matrix error (*ϵ* = 0.014). This is only slightly higher than ConDoR* in mutation matrix error (*ϵ* = 0.013) but substantially outperforming ConDoR* in clone lifespan error (*δ* = 0.091). In contrast, COMPASS has substantially higher clone lifespan error (*δ* = 0.167) and mutation matrix error (*ϵ* = 0.028). Both ConDoR* and COMPASS infer phylogenies with many longitudinal violations, while Phyllochron explicitly models longitudinal constraints and thus has no longitudinal violations even in the presence of observational uncertainty (Figure 2(E)). Notably, ConDoR* achieves low mutation matrix error (Figure 2(F)), but fails to accurately infer clone lifespans, suggesting possible overfitting and indicating that the mutation matrix error alone is an insufficient metric for assessing the quality of mutation inference. In contrast, Phyllochron has comparable mutation matrix error to ConDoR* while achieving substantially lower clone lifespan error.

### 2.4 Longitudinal phylogeny of an Acute Myeloid Leukemia patient

We used Phyllochron to analyze longitudinal single-cell DNA sequencing data from an acute myeloid leukemia (AML) patient AML-99 from [36] and compared the results with those from a state-of-the-art tumor phylogeny inference method COMPASS. This data consists of *k* = 5 longitudinal samples of the same patient taken over approximately one year. The first sample (*t*_1_) was taken at the time of diagnosis, followed by two samples, *t*_2_ and *t*_3_, taken after 2 and 6 cycles of azacitidine and enasidenib therapy, respectively. The patient was then sampled (*t*_4_) after relapsing one year later, which was followed by 18 cycles of the azacitidine and enasidenib therapy. The final sample (*t*_5_) was taken after the patient switched to salvage chemotherapy with cladribine, low-dose cytarabine, enasidenib, and venetoclax. Targeted single-cell DNA sequencing was performed for each sample using the Mission Bio Tapestri platform to measure mutations in 37 frequently mutated genes in AML using a panel of 279 amplicons (median length 216 bps) [36]. In total, this data contains a total of 19300 cells from these 5 samples (median 3992 cells per sample), and we identified 11 mutations of interest in driver genes across these cells (details in Supplementary Section D and Supplementary Section E). For this comparison, we use the COMPASS model of observed read counts in Phyllochron so that differences between the two methods are not due to differences in the observation model – further details are in Supplementary Section B.5. We also inferred a Phyllochron tree using the ConDoR read count error model (Supplementary Figure S5). We set the clone prevalence threshold (Supplementary Section B.2) to *z* = 0.001 to avoid trivially longitudinal phylogenies (Supplementary Section B.1).

Phyllochron reconstructs a longitudinally consistent phylogeny, revealing the presence of multiple cancer clones during the cycles of azacitidine and enasidenib therapy (*t*_2_, *t*_3_) which were missed by existing methods. Phyllochron infers the presence of 11 cancer clones, which we denote with labels C0 to C10 (Figure 3(A)). COMPASS (Figure 3(B)) reconstructs the same set of clones as Phyllochron, but the presence and absence of these clones at several timepoints differs from Phyllochron and contains longitudinal violations. Specifically, the COMPASS phylogeny contains multiple instances of clones disappearing at one timepoint and subsequently reappearing at a later timepoint. Clones C2 and C6, marked by the addition of mutations IDH2 p.R140Q and IDH1 p.132 respectively, disappear at *t*_3_ and reappear at *t*_4_; clone C9 marked by the gain of mutation NRAS C/T disappears at *t*_2_ and reappears at *t*_3_. Such non-contiguous clonal lifespans are not observed in the Phyllochron phylogeny. Moreover, the COMPASS phylogeny also contains instances of clones appearing before their ancestors. Specifically, clone C7, characterized by mutation RUNX1 A/G, first appears at *t*_3_, even though its descendant clone C8 is present at earlier timepoints *t*_1_ and *t*_2_. In contrast, Phyllochron finds evidence of 8 cells from clone C7 present at timepoints *t*_1_ and *t*_2_. In total, Phyllochron assigns 23 cells to different clones compared with the COMPASS solution, with 14 of the reassigned cells removing the longitudinal violations observed in the COMPASS phylogeny. Thus, the Phyllochron tree reveals that clones C6, C7, C9, and C10 persist at low prevalence at timepoints where COMPASS fails to detect them.

**Fig. 3.**
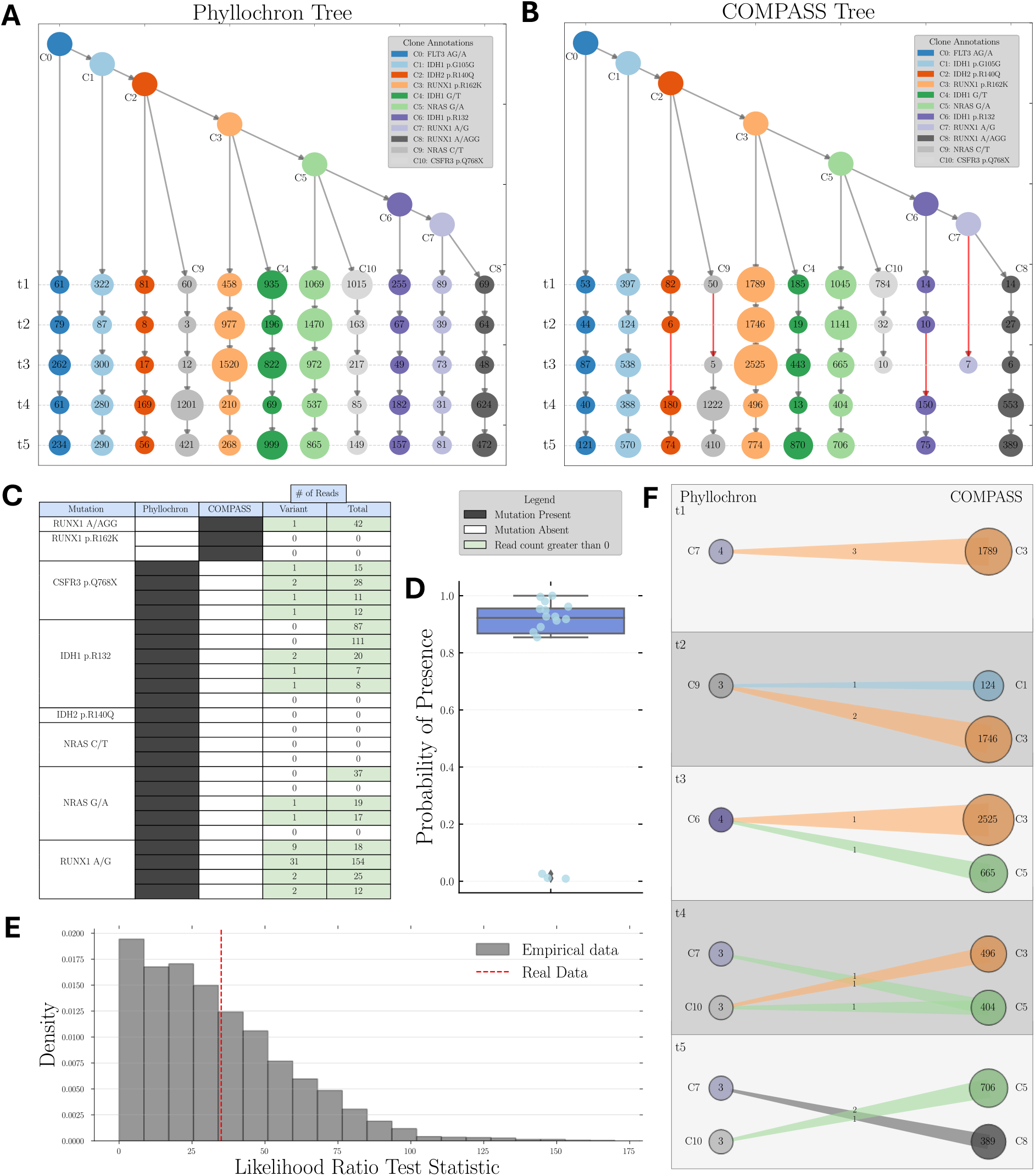
Phyllochron infers a longitudinal perfect phylogeny from longitudinal targeted single-cell sequencing data of an acute myeloid leukemia patient. (A) Phylogeny and cell assignments inferred by Phyllochron. Circles at the timepoints *t*_1_,…, *t*_5_ indicate the presence of the corresponding clone at that timepoint and numbers inside each circle indicate the number of cells assigned to the clone. (B) Phylogeny and cell assignments inferred by COMPASS. Red arrows indicate a longitudinal violation. (C) 26 loci (entries of the mutation matrix) that differ in the mutation matrices imputed by Phyllochron and COMPASS. Each row of corresponds to a mutation in a particular cell (the same mutation may differ in multiple cells), and columns indicate the presence/absence of the mutation in each method’s mutation matrix as well as the number of variant and total reads for the mutation in the cell. (D) The probability of presence under the beta-binomial read count error model for 17 loci from (C) with total read count *r* > 0. Average probability of presence is 0.76. Box plot shows the median and the interquartile range (IQR), and the whiskers denote the lowest and highest values within 1.5 times the IQR from the first and third quartiles, respectively. (E) The empirical distribution of the LRT statistic for *B* = 5000 samples constructed from the Phyllochron hypothesis test. The red dashed line represents the LRT statistic *λ*_real_ = 35 for patient AML-99. (F) The clones with different number of cells assigned by Phyllochron and COMPASS for all cells with differing assignments between the two trees. The number in the circle indicates the number of cells assigned to a clone at each timepoint.

The read count data support the mutations imputed by Phyllochron to enforce longitudinal consistency of the clone lifespans. Specifically, we examined the read counts at all 26 loci where COMPASS and Phyllochron disagree on the inferred presence or absence of mutations. Of the 3 loci inferred as absent by Phyllochron, none had more than one variant read. Among the 23 inferred as present, 13 had at least one variant read and 7 had zero variant reads (Figure 3(C)). Of the 16 mutations containing at least one total read that Phyllochron infers as present but COMPASS infers as absent, the average probability that these mutations are present under the beta-binomial model is 0.76, providing strong support for the presence of these mutations and Phyllochron’s placement of these cells into clones (Figure 3(D)).

Finally, we evaluated the evidence for a longitudinal phylogeny on the AML-99 dataset using the Phyllochron hypothesis testing procedure. We fail to reject the null hypothesis that a longitudinal phylogeny generated this dataset (Figure 3(E)), obtaining a likelihood ratio test (LRT) statistic *λ* = 35.00, corresponding to a one-sided p-value of *p* = 0.405 using an empirical distribution with *B* = 5000 samples. This suggests that the longitudinal violations observed in the COMPASS tree are not strongly supported by the data.

We further examined the 14 cells with longitudinal violations in the COMPASS solution that were reassigned to different clones by Phyllochron. In every case, including at multiple timepoints, the cells are reassigned from highly prevalent clones with large populations in the COMPASS tree to rare clones at the same timepoint in the Phyllochron tree (Figure 3(F)). We argue that the non-uniform clone attachment prior in COMPASS biases the imputation of mutations in the low confidence loci in these cells in order to assign them to highly prevalent clones. In contrast, Phyllochron does not have a bias toward assigning cells to larger clones and instead imputes mutations to avoid longitudinal violations. Thus, the longitudinal constraints facilitate the accurate recovery of rare clones that might otherwise be erroneously grouped in larger clones with similar mutation profiles.

## 3 Discussion

We introduce Phyllochron, an algorithm to infer phylogenetic trees from single-cell longitudinal DNA sequencing data. Phyllochron is based on the *longitudinal perfect phylogeny* model, a generalization of the perfect phylogeny model [44], that we introduce to describe cases where cells are measured at multiple timepoints during an evolutionary process. Phyllochron imputes a mutation matrix conforming to a longitudinal perfect phylogeny using a probabilistic model for read counts. We also introduce a hypothesis test to evaluate whether observed read counts of mutations support a longitudinal perfect phylogeny. We show that Phyllochron outperforms state-of-the-art tumor phylogeny inference methods on simulated data. On longitudinal targeted single-cell DNA sequencing data from acute myeloid leukemia (AML) patients, Phyllochron infers tumor phylogenies that are more biologically plausible than phylogenies inferred by existing methods, providing important insights regarding the persistence of rare cancer clones during treatment.

There are several limitations and directions for future research. First, in this work, we focused on single nucleotide variants (SNVs) as phylogenetic markers of tumor evolution and used the perfect phylogeny model to describe their evolution. Extending the model to other types of mutations and evolutionary models would be valuable. In particular, copy number aberrations (CNAs) are common in cancer and several single-cell sequencing technologies are tailored to measure CNAs in tumors [47–49]. An extension of Phyllochron to incorporate CNAs into longitudinal phylogenies would be valuable. CNAs require more complex models to describe the evolution of SNVs such as the *k*-Dollo [45], finite-sites [50] or constrained *k*-Dollo [25] models. Second, Phyllochron currently models the lifespan of clones with discrete timepoints. Extending the method using a continuous-time Markov model [51] would allow simultaneous inference of the evolution, clonal composition and mutation rates in the tumor. Lastly, while we focus on tumor phylogeny inference and the perfect phylogeny model, our framework for longitudinal phylogenies is not specific to cancer. For example, longitudinal sampling is also used for lineage tracing in developmental systems [52], where our algorithm could be applied with lineage tracing evolutionary models [53, 54] to reconstruct developmental lineage trees from longitudinal data. As longitudinal single-cell sequencing becomes more commonplace in cancer, developmental, and other biological studies, Phyllochron provides a foundation for future development of models and algorithms for accurate phylogeny inference from longitudinal sequencing data.

## 4 Methods

Longitudinal single-cell sequencing experiments track tumor evolution by profiling cells at multiple timepoints. Specifically, longitudinal scDNA-seq data is represented by a binary mutation matrix *A* ∈ {0, 1}^*n*×*m*^, where *a*_*i,j*_ = 1 indicates that mutation *j* is present in cell *i*, and a timepoint vector *τ*, where *τ*_*i*_ ∈ {1,…, *k*} denotes the timepoint when cell *i* was sampled. To model the generative process behind the observed mutation information *A* and temporal information *τ*, we introduce longitudinal phylogenies derived from constraints imposed by longitudinal sampling.

### 4.1 Longitudinal Phylogenies

We formally define a longitudinal phylogeny for a mutation matrix *A* and timepoints ***τ*** following the temporal consistency property and the ancestral sampling assumption as follows.

#### Longitudinal phylogeny

*A* longitudinal phylogeny *for a mutation matrix A and timepoints* ***τ*** *is a rooted tree T in which the leaves and some subset of the internal vertices are labeled by rows* **a**_*i*_ *of A and timepoints τ*_*i*_ ∈ {1,…, *k*} *such that for every path P from root to leaf, the timepoint labels τ*_*i*_ *on P appear in consecutive and strictly increasing order* (1,…, *k*′) *for some* 1 ≤ *k*′ ≤ *k*.

Longitudinal phylogenies are distinct from timed phylogenies, which have been previously used to model temporal sequencing data [41–43]. Timed phylogenies are trees in which the sequenced cells correspond to leaves and the internal vertices represent ancestral cells. The branches of a timed phylogeny have lengths such that the measurement time of a cell is given by the total length of the path from the root to the leaf corresponding to that cell. In a longitudinal phylogeny, we represent some of the ancestral cells by sequenced cells that have identical mutational profiles. This allows us to impose temporal dependence across successive longitudinal measurements.

### 4.2 Longitudinal Perfect Phylogenies

Constructing a longitudinal phylogeny from sequencing data requires an evolutionary model that describes the allowed mutation state transitions along the edges of the phylogeny. The simplest model used in cancer genomics is the *perfect phylogeny* model [44], where each mutation locus in a cell has one of two states: state 0 representing the absence of the mutation, and state 1 representing the presence of the mutation. In a perfect phylogeny, the state of each locus can transition from the unmutated state (0) to the mutated state (1) at most once, and cannot subsequently transition back to the unmutated state.

Given a mutation matrix *A*, a phylogeny *T* is a perfect phylogeny for *A* if there exists a way to assign each mutation to exactly one edge of *T* such that mutation *j* labels an edge on the path from root to the vertex corresponding to cell *i* if and only if *a*_*i,j*_ = 1. Note that not all mutation matrices will admit a perfect phylogeny and we call *A* a perfect phylogeny matrix if and only if there exists a perfect phylogeny *T* for *A*. We define a *longitudinal perfect phylogeny* to be a longitudinal phylogeny where the mutations satisfy a perfect phylogeny.

#### Longitudinal Perfect Phylogeny

*A longitudinal phylogeny T for mutation matrix A and timepoints* ***τ*** *is a* longitudinal perfect phylogeny *provided T is a perfect phylogeny for A*.

The longitudinal perfect phylogeny model is a generalization of the perfect phylogeny model, which pertains to the special case where the number of timepoints *k* = 1, i.e. when each cell *i* is sequenced at the same timepoint *τ*_*i*_ = 1. Below we state three problems for longitudinal perfect phylogenies which generalize problems that are well-studied for perfect phylogenies.

First, while there is a well-known characterization of perfect phylogeny matrices, i.e. mutation matrices that admit a perfect phylogeny [44, 55] (Theorem 4.1 below). Not all pairs of perfect phylogeny matrices and timepoint vectors will admit a longitudinal perfect phylogeny. This motivates the following problem.

#### Longitudinal Perfect Phylogeny (LPP)

*Given a mutation matrix A and timepoint vector* ***τ***, *determine if A and τ admit a longitudinal perfect phylogeny, and if so, construct one*.

Second, missing data is a common feature of mutation matrices derived from single-cell sequencing data. A mutation matrix is called *incomplete*, if it has missing entries, typically denoted by ‘?’. Given an incomplete mutation matrix*A*′ ∈ {0, 1, ?}^*n*×*m*^ a *completion* of *A*′ is a binary mutation matrix *A*∈ {0, 1}^*n*×*m*^ which is consistent with *A*′ on all the 1 and 0 entries, i.e. where *a*_*i,j*_ = 1 if 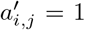 and *a*_*i,j*_ = 0 if 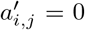. The problem of finding a *completion A* of an incomplete mutation matrix *A*′, i.e. imputing the missing entries of *A*′, that admits a perfect phylogeny has been extensively studied [56, 57]. We generalize this problem in the longitudinal context as follows.

#### Incomplete Longitudinal Perfect Phylogeny (ILPP)

*Given an incomplete mutation matrix A*′ *and timepoint vector* ***τ***, *find completion A of A*′, *if one exists, such that A and* ***τ*** *admit a longitudinal perfect phylogeny*.

Third, in practice, we do not measure the mutation matrix *A* directly. Instead, sequencing technologies yield counts of reference and variant reads at mutation loci in each sequenced cell. Specifically, let *Q* ∈ ℤ^*n*×*m*^ be the variant read count matrix, where *q*_*i,j*_ is the number of reads with the variant allele for mutation *j* in cell *i*, and let *R* ∈ ℤ^*n*×*m*^ be the total read count matrix, where *r*_*i,j*_ is the total number of reads for mutation *j* in cell *i*. Since the cells and the mutations in each cell are sequenced independently, the likelihood Pr(*Q*| *R, A*) of observing the variant read count matrix *Q* for given total read count matrix *R* and mutation matrix *A* is

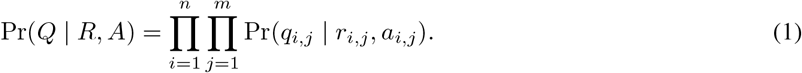

We model the probability Pr(*q*_*i,j*_ | *r*_*i,j*_, *a*_*i,j*_) of observed variant read counts *q*_*i,j*_ for an individual cell and mutation using a beta-binomial distribution, similar to previous work [58, 22, 59, 25]. Supplementary Section C.1 describes the probabilistic model for the read counts in detail.

For given read count matrices *Q* and *R* and timepoints vector ***τ***, our goal is to construct a mutation matrix *A* that maximizes the likelihood defined in Equation 1 such that *A* and *τ* admit a longitudinal perfect phylogeny *T*. We refer to this as the *Longitudinal Perfect Phylogeny from Read Counts* (LPP-RC) problem and pose it as follows.

#### Longitudinal Perfect Phylogeny from Read Counts (LPP-RC)

*Given a variant read count matrix Q, total read count matrix R, and a timepoint vector* ***τ***, *find a mutation matrix A and longitudinal perfect phylogeny T such that (i) A maximizes the likelihood* Pr(*Q* | *R, A*) *and (ii) A and* ***τ*** *admit T*.

### 4.3 Combinatorial Characterization of Longitudinal Perfect Phylogenies

We derive a combinatorial characterization of mutation matrices that admit a phylogeny under the longitudinal perfect phylogeny model and determine the computational complexity of the phylogeny inference problems described above. We build on previous work on characterization of mutation matrices that admit a phylogeny under the perfect phylogeny model, which is a special case of our model when the number *k* of timepoints is 1. Previous works [44] have shown that a mutation matrix is a perfect phylogeny matrix, i.e. it admits a perfect phylogeny, if and only if it satisfies the *three gametes condition*, which is stated as follows.

#### Theorem 4.1

(Gusfield 1991 [44]). *A binary matrix A* ∈ {0, 1}^*n*×*m*^ *admits a perfect phylogeny if and only if no two columns of A contain the three pairs* (0, 1), (1, 0) *and* (1, 1).

For a *n* × *m* mutation matrix *A*, the perfect phylogeny, if it exists, is unique and can be constructed in *O*(*nm*) time [44]. However, this perfect phylogeny may not be consistent with the timepoint vector ***τ*** under the longitudinal perfect phylogeny constraints (see Figure 4 for example). In the following, we derive additional conditions that mutation matrix *A* and timepoints ***τ*** must satisfy to admit a longitudinal perfect phylogeny when the number *k* of timepoints is greater than 1.

**Fig. 4.**
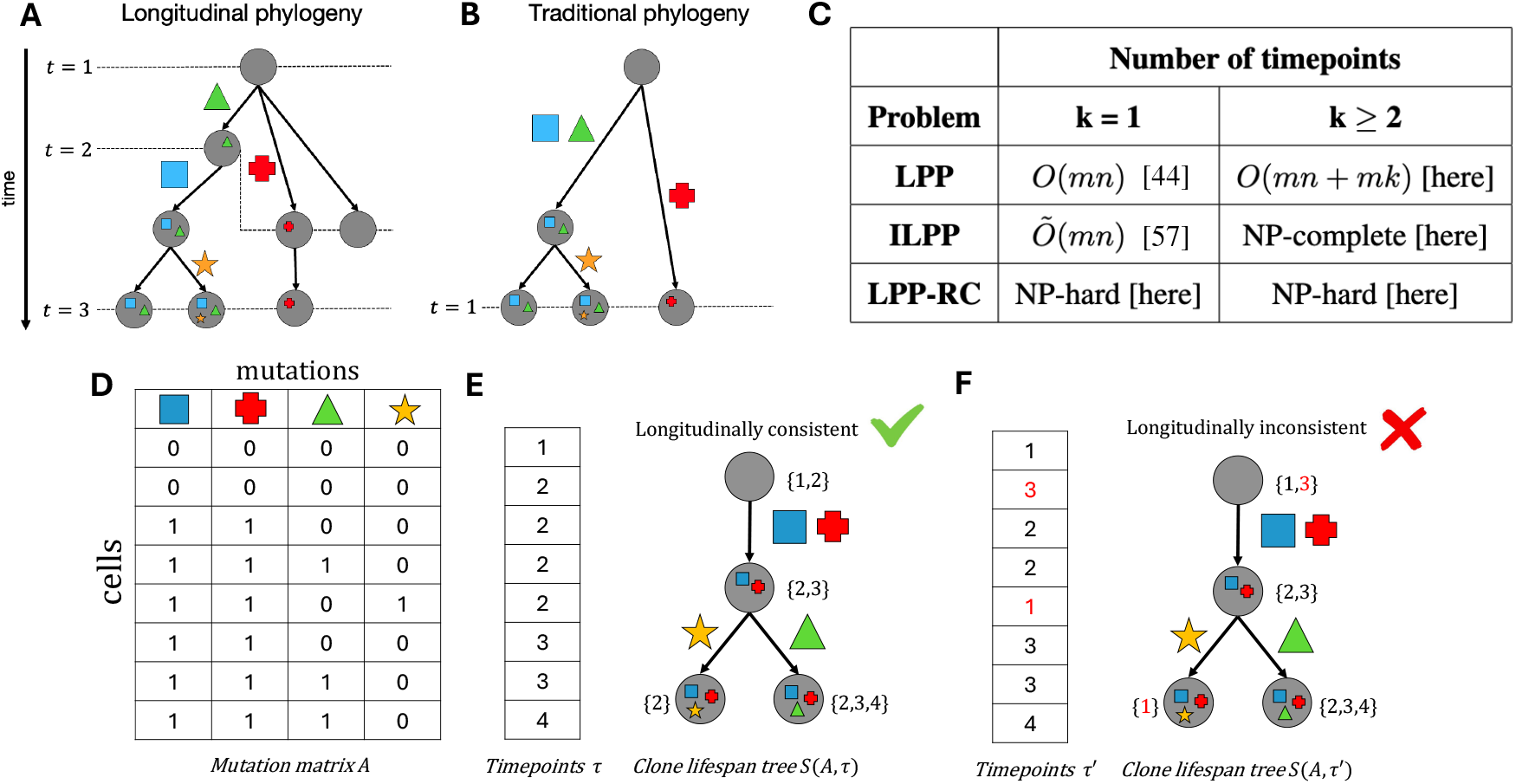
(A) Longitudinal phylogenies are rooted trees in which the leaves and some of the internal vertices represent the cells sequenced from the longitudinal samples. (B) Traditional phylogenies represent all the sequenced cells by leaves. (C) Summary of the complexity results derived in this study put in context with previous work. (D) A mutation matrix *A* that admits a perfect phylogeny. (E) Timepoints vector *τ* admits a longitudinally consistent clone lifespan tree *S*(*A, τ*) with *A*. (F) Timepoint vector *τ*′ yields a longitudinally inconsistent clone lifespan tree *S*(*A, τ*′).

We introduce the *clone lifespan tree* to represent the evolution and lifespan of clones in the tumor. A clone is a subpopulation of cells with the same set of mutations, which correspond to a set of identical rows in the mutation matrix *A*. Since a perfect phylogeny matrix *A* with *m* mutations can have at most *m* distinct rows and the number *n* of cells is typically much larger than *m*, duplicate rows in *A* are often observed in real data. Let *σ* be a partition of cells into cancer clones such that *σ*(*i*) indicates the clone of cell *i*. The vertices of a clone lifespan tree represent cancer clones and edges represent ancestral relationship between these clones. Each edge of the clone tree is labeled by mutations that distinguish the parent and the child clone, and satisfy the following two conditions. First, each mutation labels exactly one edge of the tree. Second, a mutation appears on the path from root to a clone if and only if cells of that clone contain that mutation. Each clone, i.e. vertex of the tree, is labeled by its *lifespan*, which is the set of timepoint where it was observed in the tumor. Specifically, lifespan of clone *u* is *ℓ* (*u*) = {*τ*_*i*_ : *i* ∈ *σ*^−1^(*u*)}. The clone lifespan tree without the lifespan labels is referred to as *clone tree* [60, 61, 27]. We show that for a perfect phylogeny matrix *A* and timepoints vector ***τ***, the clone lifespan tree *S*(*A*, ***τ***) is unique.

#### Lemma 4.1.

*The clone lifespan tree S*(*A*, ***τ***) *for a perfect phylogeny matrix A and timepoints* ***τ*** *is unique*.

The lifespans of the clones must satisfy certain constraints to be *longitudinally consistent*. First, a clone must be observed for a contiguous time interval in the tumor, i.e. the lifespan must be a contiguous set of consecutive timepoints. Specifically, if a clone is not observed at some timepoint, it can not reappear at a later timepoint. Second, the time at which a clone arises in a tumor, its ancestor must also be present. Let *π*(*v*) be the nearest ancestor of clone *v* in *S*(*A*, ***τ***) such that *ℓ*(*π*(*v*)) ≠ ∅. We require that the lifespan of *π*(*v*) must overlap with the lifespan of *v*. We formally state these conditions to characterize mutation matrices *A* and timepoint vectors ***τ*** that admit a longitudinal perfect phylogeny in the following theorem.

#### Theorem 4.2.

*Mutation matrix A and timepoint vector* ***τ*** *admit a longitudinal perfect phylogeny if and only if A is a perfect phylogeny matrix and the clone lifespan tree S*(*A*, ***τ***) *is longitudinally consistent, i*.*e. it has the following properties*

1. *The lifespan ℓ*(*u*) *of each clone is a contiguous set, i*.*e. each timepoint p in the interval* [min(*ℓ*(*u*)), max(*ℓ*(*u*))] *is present in ℓ*(*u*).
2. *For each clone v, we have* min(*ℓ*(*π*(*v*))) ≤ min(*ℓ*(*v*)) ≤ max(*ℓ*(*π*(*v*))).

Using this characterization, we show that a longitudinal phylogeny for *A* and ***τ***, if it exists, can be constructed in polynomial time.

#### Theorem 4.3.

*Given a mutation matrix A* ∈ {0, 1}^*n*×*m*^ *and timepoints vector τ* ∈ {1,…, *k*}^*n*^, *a longitudinal perfect phylogeny for A and τ can be constructed in O*(*nm* + *mk*) *time, if one exists*.

In Supplementary Sections A.4 and A.5, we prove the hardness of the Incomplete Longitudinal Perfect Phylogeny (ILPP) problem 4.2 and the Longitudinal Perfect Phylogeny from Read Counts (LPP-RC) problem 4.2, respectively.

### 4.4 Phyllochron: longitudinal phylogeny inference algorithm

The longitudinal phylogeny inference algorithm consists of two main components: (i) a mixed integer linear program (MILP) that leverages the characterization of a longitudinal perfect phylogeny (Section 4.3) to assign cells to a fixed clone tree *S*, ensuring that the assignment is longitudinal and maximizes the likelihood of the read count data under our model Pr(*Q* | *R, A*); and (ii) a tree search procedure that explores the space of clone trees using Subtree-Pruning-Regrafting (SPR) [62–65] moves (Section 4.4.2). We alternate between these two steps until the algorithm converges to a local optimum in both tree structure and cell assignment.

#### 4.4.1 Phyllochron MILP formulation

Here, we describe the MILP used in Phyllochron to build longitudinal phylogenies for a fixed clone tree. Let *S* be the clone tree over 𝒞, where 𝒞 is the set of all clones. We introduce binary variables *x*_*i,c*_ to indicate if cell *i* is assigned to clone *c* ∈ 𝒞. The lifespan label *ℓ*(*c*) of a clone *c* is encoded by continuous variables *τ*_*c,t*_ ∈ [0, 1] for *t* ∈ {1,…, *k*} such that *τ*_*c,t*_ = 1 if and only if *t* ∈ *ℓ*(*c*).

##### Cell assignment constraints

Each cell must be assigned to exactly one clone, which we enforce using the following constraint,

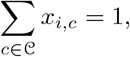

for each cell *i* ∈ {1,…, *n*}. Further, *t* ∈ {1,…, *k*} is in the lifespan *ℓ*(*c*) if and only if at least one cell *i* with timepoint *σ*(*i*) = *t* is assigned to clone *c*, which we enforce as follows.

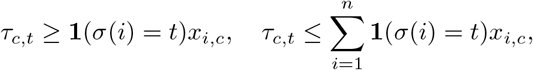

for each cell *i*, timepoint *t* and clone *c*, where **1** is the indicator function.

##### Longitudinal consistency constraints

We require that the lifespans of clones must satisfy the constraints described in Theorem 4.2. We enforce that the lifespan of each clone is a contiguous set (condition 1 in Theorem 4.2) by requiring that for any three timepoints *t*_1_ < *t*_2_ < *t*_3_, we must not observe 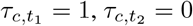 and 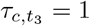, i.e.

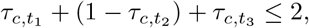

for 1 ≤ *t*_1_ < *t*_2_ < *t*_3_ ≤ *k*. According to condition 2 in Theorem 4.2, we require that: (i) min(*ℓ*(*π*(*c*))) ≤ min(*ℓ*(*c*)) and (ii) min(*ℓ*(*c*)) ≤ max(*ℓ*(*π*(*c*))) for each clone *c*. We enforce (i) by requiring that there are no pair of timepoints *t*_1_ < *t*_2_ such that clone *c* is present at time *t*_1_, i.e. 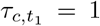, and min(*ℓ*(*π*(*c*))) = *t*_2_, i.e. 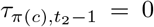 and 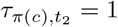. This is achieved with the following constraints,

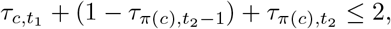

for 1 ≤ *t*_1_ < *t*_2_ ≤ *k*. Following an analogous approach, we enforce min(*ℓ*(*c*)) ≤ max(*ℓ*(*π*(*c*))). Specifically, we enforce that there must not be two timepoints *t*_1_ < *t*_2_ such that clone *π*(*c*) disappears at *t*_1_, i.e. 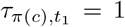 and 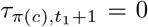, and clone *c* appears at time *t*_2_, i.e. 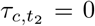 and 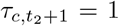. This is achieved using the following constraints for 1 ≤ *t*_1_ < *t*_2_ < *k*,

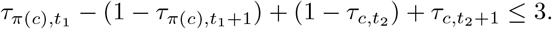

##### Unobserved clone constraints

We may not observe all ancestral clones present in 𝒞, as sets of mutations may be gained in quick succession, preventing the observation of these intermediate clones that contain subsets of these mutations. We introduce a binary variable Ω_*c*_, ∀*c* ∈ 𝒞 that indicates if clone *c* is present at any timepoint. We enforce this as follows:

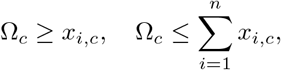

for each cell *i* and clone *c*. Next, we introduce a binary variable *γ*_*c,c*_ that indicates if clone *c*′ = *π*(*c*) is the most recent observed ancestor clone of *c*. As ancestral clones may be unobserved, *c*′ is not necessarily the most recent ancestor of *c* in *S*. We enforce this constraint as follows:

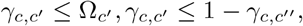

where *c*′ is an ancestor of *c* and *c*^′′^ is an ancestor of *c*′ in *S*. We add the following additional constraints:

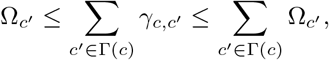

where Γ(*c*) is the ordered set of all ancestral clones of *c* where the first clone in Γ(*c*) is the root and the last clone is direct ancestor of *c*. This constraint enforces the presence of a most recent observed ancestor clone for all *c* where an ancestor *c*′ is present. Furthermore, we add the constraint 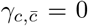 for all 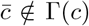. We use 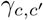 as an indicator variable that enforces longitudinal consistency constraints for all clone-ancestor pairs (*c, c*′) where 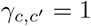.

##### Objective Function

We maximize Pr(*Q*| *R, A*) where *A* is the longitudinal perfect phylogeny mutation matrix. Let *a*_*c,j*_ denote the state of mutation *j* in clone *c* in clone tree *S*. The log-likelihood is a weighted sum of the clone assignment variables *x*_*i,c*_, where each weight *w*_*i,c*_ is determined by the likelihood of the observed total and variant read counts *r*_*i,j*_, *q*_*i,j*_ respectively when cell *i* is assigned to clone *c*, i.e. *a*_*i,j*_ = *a*_*c,j*_ for all mutations *j*.

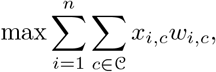

where

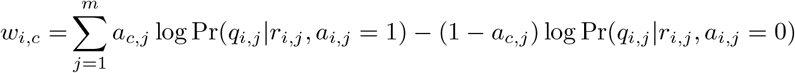

This MILP has *O*(*nm*) binary variables, *O*(*mk*) continuous variables and *O*(*nmk* + *mk*^3^) constraints. More details on the Phyllochron MILP can be found in Supplementary Section B.

#### 4.4.2 Phyllochron Tree search

The MILP introduced in Section 4.4 infers a longitudinally consistent lifespan labeling on the fixed clone tree *T*. To infer both the clone tree and the lifespan labeling, we search through the space of all possible clone trees. A brute-force search through the entire tree space would scale exponentially in the number of mutations *m*, and thus a heuristic must be employed to efficiently search through candidate tree topologies. We implement Subtree-Pruning-Regrafting (SPR) moves [66] to traverse tree space. Specifically, given a tree *T*, an SPR move prunes a subtree rooted at mutation 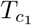, and regrafts it as the child of another mutation *c*_2_ on the original tree. Given a clone tree *T* with *m* mutations, there are *O*(*m*^2^) candidate trees generated from a single SPR move on *T*. We explore the 1-SPR move neighborhood and greedily select the clone tree with the highest likelihood using the MILP described in Section 4.4, and keep exploring this neighborhood until we reach a local optimum. In this study, we initialize Phyllochron with the clone tree *T*_0_ inferred by COMPASS [27].

### 4.5 Phyllochron: Ancestral sampling hypothesis test

To test the support for longitudinal phylogenies on our data, we compare two nested models, the longitudinal perfect phylogeny model and the perfect phylogeny model, employing a likelihood ratio test to assess whether the less restrictive model (perfect phylogeny) provides a significantly better fit to the data compared to the restrictive one (longitudinal perfect phylogeny) given a clone tree *T*.

Let ***PP*** represent the set of perfect phylogeny mutation matrices on *T* and let ***LP P*** represent the set of longitudinal perfect phylogeny mutation matrices on *T*. Clearly, ***LPP*** ⊂ ***PP***. We define the null and alternative hypotheses as follows:

- **Null Hypothesis** (*H*_0_): The variant read counts *Q* are generated by *A* ∈ ***LP P***.
- **Alternative Hypothesis** (*H*_*A*_): The variant read counts *Q* are generated by *A* ∈ ***PP***.

Let *Â*_LPP_ represent the maximum likelihood mutation matrix from Phyllochron and let *Â*_PP_ represent the maximum likelihood mutation matrix under the perfect phylogeny model. Furthermore, let *L*(*A*) be the probability of observing the variant read counts 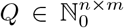 given total read counts 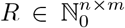 and mutation matrix *A*. That is, variant read counts *Q*_1,1_,…, *Q*_*n,m*_ are i.i.d. with likelihood 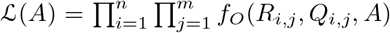 where *f*_*O*_(*R*_*i,j*_, *Q*_*i,j*_) is the probability mass/density function under any read count model *O*. The likelihood ratio test (LRT) statistic is thus defined as:

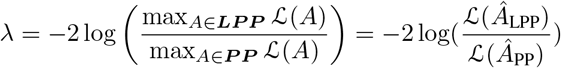

We compute a one-sided p-value using the empirical distribution of the LRT statistic *λ*, as follows.

Given a clone tree *T*, total read counts *R*, some read count model *O* and clone sizes *k*_*m,t*_ corresponding to the number of cells assigned to clone *m* at timepoint *t* in the maximum likelihood mutation matrix *Â*_LPP_, we generate sample log likelihood ratios *λ*_1_,…, *λ*_*D*_ using the following procedure.

1. **Sample cell assignments:** We generate mutation matrices *A*^*i*^ by randomly sampling the clone assignment of cells at each timepoint *t* without replacement such that 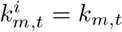.
2. **Generate variant read counts:** Given the total read counts *R* and sampled mutation matrix *A*^*i*^, generate the variant read counts *Q* by sampling from the probability mass function of the underlying read count model *O*. This produces a variant read count dataset *Q*^*i*^.
3. **Compute the likelihood ratio test statistic:** Compute 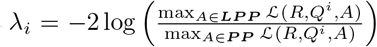, where likelihood function ℒ depends on the underlying read count model *O*. Under the beta-binomial read count model, the numerator is equivalent to the inferred Phyllochron matrix on fixed-tree *T* and the denominator is equivalent to the inferred ConDoR matrix on *T*.

### 4.6 Simulation performance metrics

To evaluate the accuracy of the inferred phylogenies and mutation profiles on simulated data, we define quantitative metrics that compare the inferred results to the known ground truth. These metrics allow us to assess both the topological similarity of the trees and the accuracy of the inferred cell-to-clone assignments.

We use three metrics, the clone lifespan error, the mutation matrix error and number of longitudinal violations, to evaluate the performance of competing methods on simulated data. The clone lifespan error quantifies the accuracy of the clone lifespans inferred by each method. Let 𝒞 be the set of simulated clones, *ℓ*(*c*) denote the lifespan of clone *c* and 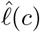 be the lifespan of clone *c* inferred by a method. The clone lifespan error 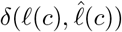 is the average Jaccard distance between *ℓ*(*c*) and 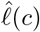 across all clones *c* ∈ 𝒞 given by

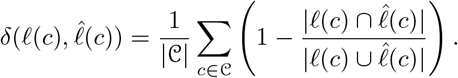

The mutation matrix error quantifies the accuracy of the mutational profiles of the cells inferred by each method. The mutation matrix error *ϵ*(*A, Â*) between a ground-truth mutation matrix *A* and inferred mutation matrix *Â* is the normalized L_1_ norm between *A* and *Â* given by

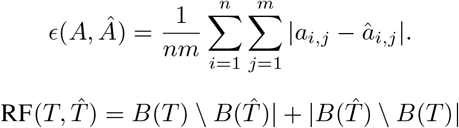

where *B*(*T*) represents the set of bipartitions uniquely defining *T*.

The number of longitudinal violations quantifies the compatibility of inferred cell-to-clone assignments under our longitudinal framework, and thus we define a longitudinal violation score that quantifies violations of the temporal consistency and ancestral sampling assumption for inferred phylogenies.

Let *A* be the inferred *n* × *m* binary mutation matrix and let ***τ*** be the timepoint vector encoding the sampling time *τ*_*i*_ for each cell *i*. Each unique mutation profile (row of *A*) defines a putative inferred clone, and we associate each clone with the multiset of timepoints at which its corresponding cells were observed.

We identify two types of violations:

#### Temporal inconsistency

A clone is expected to appear in a temporally contiguous interval of timepoints, reflecting uninterrupted clonal persistence over time. For each inferred clone, we extract its lifespan *τ* (*c*) *⊆* 1,…, *k*, the set of timepoints where it is observed. If *τ* (*c*) is not a complete interval (i.e., contains gaps), we record a temporal consistency violation. This captures violations of the requirement that labeled vertices along a path in a longitudinal phylogeny must increase consecutively in time.

#### Ancestral sampling violation

Under the ancestral sampling assumption, each sampled clone at time *t*> 1 must be the descendant of a sampled clone present at time *t* − 1. To test this, we define a parent-child relationship between clones based on inclusion of mutation profiles (i.e., candidate parent clones are those with a subset of the child clone’s mutations) as per the perfect phylogeny constraints. For each inferred clone *c*, we identify the parent clone *c*′ and verify that min *τ* (*c*′) < min *τ* (*c*) and max *τ* (*c*′) ≥ min *τ* (*c*), as required for parent clones to precede or temporally overlap with their descendants. A violation is recorded if either condition fails.

The total longitudinal violation score is the number of inferred clones that violate at least one of these criteria. The first two evaluation metrics are 0 if and only if the inferred results match the ground-truth. The last evaluation metric is 0 if and only if the inferred results are longitudinal.

## 5 Software Availability

Phyllochron is available on Github (https://github.com/raphael-group/Phyllochron).

## 7 Acknowledgments

This work was supported by NIH/NCI grants U24CA264027 and U24CA248453 awarded to B.J.R.

## Footnotes

1 ConDoR* indicates Phyllochron without longitudinal constraints, which is equivalent in performance to the original ConDoR.

